# Antigen structure affects cellular routing through DC-SIGN

**DOI:** 10.1101/566141

**Authors:** Cassie M. Jarvis, Daniel B. Zwick, Joseph C. Grim, Lynne R. Prost, Mohammad Murshid Alam, Jaye C. Gardiner, Soyeong Park, Laraine L. Zimdars, Nathan M. Sherer, Laura L. Kiessling

## Abstract

Dendritic cell (DC) lectins mediate the recognition, uptake, and processing of antigens, but they can also be co-opted by pathogens for infection. These distinct activities depend upon the routing of antigens within the cell. Antigens directed to endosomal compartments are degraded and the peptides presented on MHC class II molecules thereby promoting immunity. Alternatively, HIV-1 can avoid degradation, as virus engagement with C-type lectin receptors (CLRs), such as DC-SIGN, results in trafficking to surface-accessible invaginated pockets. This process appears to enable infection of T cells in *trans*. We sought to explore whether antigen fate upon CLR-mediated internalization was affected by antigen physical properties. To this end, we employed the ring-opening metathesis polymerization to generate glycopolymers that each display multiple copies of mannoside ligand for DC-SIGN yet differ in length and size. The rate and extent of glycopolymer internalization depended upon polymer structure—longer polymers were internalized more rapidly and more efficiently than were shorter polymers. The trafficking, however, did not differ, and both short and longer polymers colocalized with transferrin-labeled early endosomes. To explore how DC-SIGN directs larger particles, such as pathogens, we induced aggregation of the polymers to access particulate antigens. Strikingly, these particulate antigens were diverted to the invaginated pockets that harbor HIV-1. Thus, antigen structure has a dramatic effect on DC-SIGN-mediated uptake and trafficking. These findings have consequences for the design of synthetic vaccines. Additionally, the results suggest new strategies for targeting DC reservoirs that harbor viral pathogens.

**Significance Statement:** Dendritic cells (DCs) express cell-surface proteins (lectins) that bind to carbohydrates displayed on the surface of pathogens. The binding of pathogens to these lectins results in internalization to endosomal compartments where the pathogens are destroyed and an immune response is initiated. HIV-1 can subvert this process – lectin engagement routes HIV-1 to cellular compartments that allow the virus to evade destruction. We synthesized glycopolymers to test whether the size and physical properties of the antigens impact trafficking. Small polymers trafficked to endosomes, as expected. Alternatively, large particulate polymers localized in the non-endosomal compartments occupied by HIV. These data indicate that antigen structure affects routing by DC lectins. Our findings can be exploited to direct cargo to different compartments.

## Introduction

Dendritic cells (DCs) are sentinels of the innate immune system: they sense pathogens in the periphery and mount adaptive responses to enable immunity. Glycans are key determinants of pathogen recognition by DCs. Their importance is reflected in the multiplicity of carbohydrate-binding protein (lectin) receptors present on the surface of DCs (1, 2). Lectins can recognize antigenic glycans to facilitate pathogen internalization and subcellular trafficking to endosomal compartments (3); pathogens are then degraded and antigenic peptides are loaded onto major histocompatibility complexes (MHC) for presentation to T cells.

Despite the apparent function of lectins in mediating immunity, viruses have developed mechanisms to co-opt lectins for dissemination (4–6). The complicated relationship between HIV-1 and the lectin DC-SIGN (dendritic cell-specific ICAM-3-grabbing non-integrin) serves as an example. DC-SIGN is a member of the family of Ca^2+^-dependent lectin receptors (C-type lectin receptors or CLRs) that recognizes high mannose- and fucose-substituted glycans. DC-SIGN binds mannosylated gp120 on the HIV-1 envelope to mediate internalization (7, 8). Curiously, the virus is not routed to the traditional endocytic pathway nor is it degraded; instead, it is trafficked to subcellular compartments near the cell surface (9). These “invaginated pockets”, which are enriched in tetraspanins such as CD81, can facilitate viral transfer to T cells (i.e. *trans* infection) (10, 11). Similarly, Siglec-1 (12), another DC lectin, is implicated in facilitating HIV-1 *trans* infection of T cells (13). Thus, HIV-1 exploits lectin trafficking to avoid degradation and to promote infection of CD4^+^ T cells *in trans* (7).

These studies indicate that DCs can take up glycosylated antigens and traffic them either to the endosomes or to invaginated pockets. In the case of HIV-1, recent studies implicate actin dynamics (10). Multivalent glycosylated antigens like HIV-1 can promote the clustering of immune receptors to facilitate antigen uptake as well as changes in the actin cytoskeleton (14–17). We postulated that glycosylated antigens of different size or density could vary in their ability to promote lectin-mediated uptake and perhaps even trafficking (reviewed in (18)). Though this possibility seemed feasible, we could not find examples in which antigens displaying the same epitope were trafficked to distinctly different cellular locations.

The binding of multivalent antigens to immune receptors promotes receptor clustering, and the extent of clustering can influence downstream signaling and alter actin polymerization and stabilization (19, 20). DC-SIGN is a tetrameric lectin and the binding of multivalent antigens results in their uptake. Specifically, anti-DC-SIGN antibodies that can cluster DC-SIGN were internalized (21–23). Similarly, carbohydrate-substituted dendrimers undergo internalization (24), and DC-SIGN-expressing cells appeared to exhibit a preference for taking up higher generation glycodendrimers. The role of DC-SIGN clustering in antigen internalization is not apparent however, as anti-DC-SIGN Fab fragments that presumably do not cluster the lectin are also internalized (21-23). Moreover, the role of antigen structure in trafficking was unclear.

Polymeric antigens can promote changes like receptor-capping in B cells, which depends on the actin cytoskeleton (19, 25). We therefore used the ring-opening metathesis polymerization (ROMP) to generate polymers of defined length that act as ligands for DC-SIGN. The longer polymers were internalized more efficiently than their shorter counterparts, but all of these soluble antigens were routed to endosomal compartments. To access antigens that more effectively mimic the features of pathogens, we aggregated the polymers to form large particulate antigens. Remarkably, the particulate antigens were not directed to endosomes; their trafficking paralleled that of HIV-1 in that they localized in invaginated pockets that contain CD81 (9). Thus, control over antigen structure provides access to different dendritic cell compartments. These findings suggest the intrinsic, bulk properties of antigens can direct their localization in dendritic cells. Moreover, they suggest that antigen structure can be tailored for synthetic vaccines or to target viruses subverting immune surveillance.

## Results

### Internalization of glycopolymers by DC-SIGN

Carbohydrate-substituted polymers have been used to cluster cell surface receptors, including transmembrane lectins ((19, 25, 26), reviewed in (27)). As the length of the polymer increases, so does its ability to cluster cell-surface receptors (19, 25, 26, 28); therefore, polymers of controlled length can reveal the importance of receptor clustering in a given process (18, 27), The length of glycan-substituted polymers can be controlled using modern polymerization chemistry—as was first demonstrated using ROMP (**Error! Reference source not found.**) (29). We therefore used ROMP to generate glycopolymers to probe the effects of receptor clustering on DC-SIGN-mediated internalization and trafficking.

ROMP effected by defined metal carbene catalysts is a living polymerization in which the initiation rates can exceed those of propagation (30, 31). Control over the length of the resulting polymer is achieved by altering the ratio of catalyst initiator to monomer (30, 32). We generated a panel of glycopolymers of defined lengths (DP = 10, 33, 100, 275) using a functional group-tolerant ruthenium catalyst that affords polymers of controlled length and narrow polydispersity. The polymers were functionalized to display an aryl mannoside, a ligand that we found is superior to alkylmannosides for binding to DC-SIGN. The polymers also carried a fluorophore, which was used to monitor their cellular binding, uptake, and localization. The mannoside epitope density was held constant between all four polymers (∼35 mol%), such that the effects of glycopolymer length (and thus receptor clustering) and not ligand density could be assessed.

We tested whether these glycopolymers were internalized by cells expressing DC-SIGN. To analyze the role of DC-SIGN in the absence of other mannose-binding lectins, we compared glycopolymer uptake using Raji cells (a human B cell line that does not express DC-SIGN) and a stably transfected Raji cell line that expresses DC-SIGN (Raji/DC-SIGN). Cells were treated with fluorophore-labeled glycopolymers of defined length, and internalization was assessed by confocal microscopy. The glycans were internalized only by the cells expressing DC-SIGN (Fig. 2A). Moreover, polymer internalization was dose-dependent, further supporting the role for a specific interaction with DC-SIGN (Fig. S1).

**Fig. 1.**
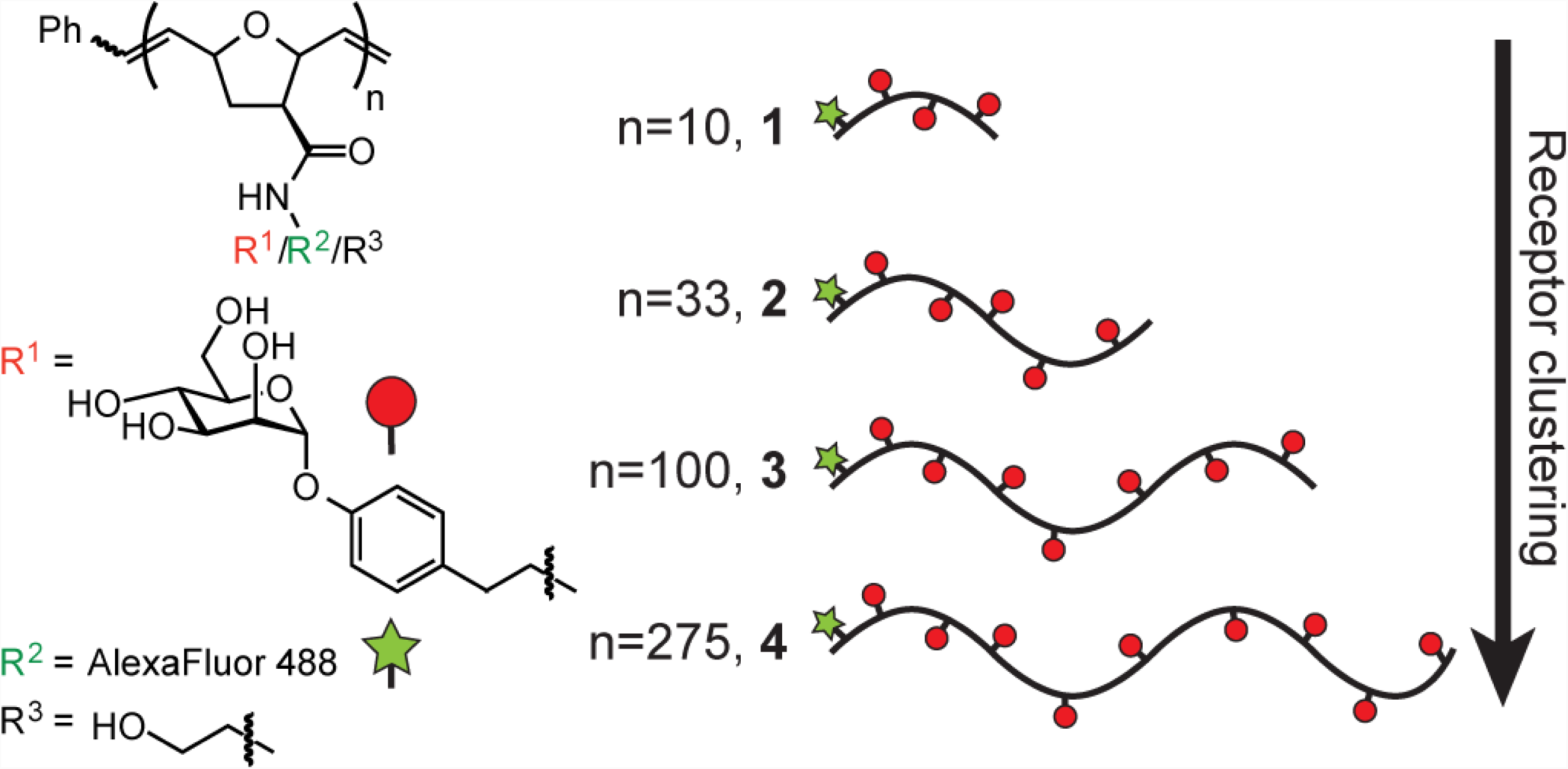
Glycopolymer probes of DC-SIGN endocytosis. A panel of glycopolymers **1** – **4** of defined length were synthesized bearing an aryl mannoside ligand (red) for DC-SIGN as well as AlexaFluor 488 (green) for visualization.

**Fig. 2.**
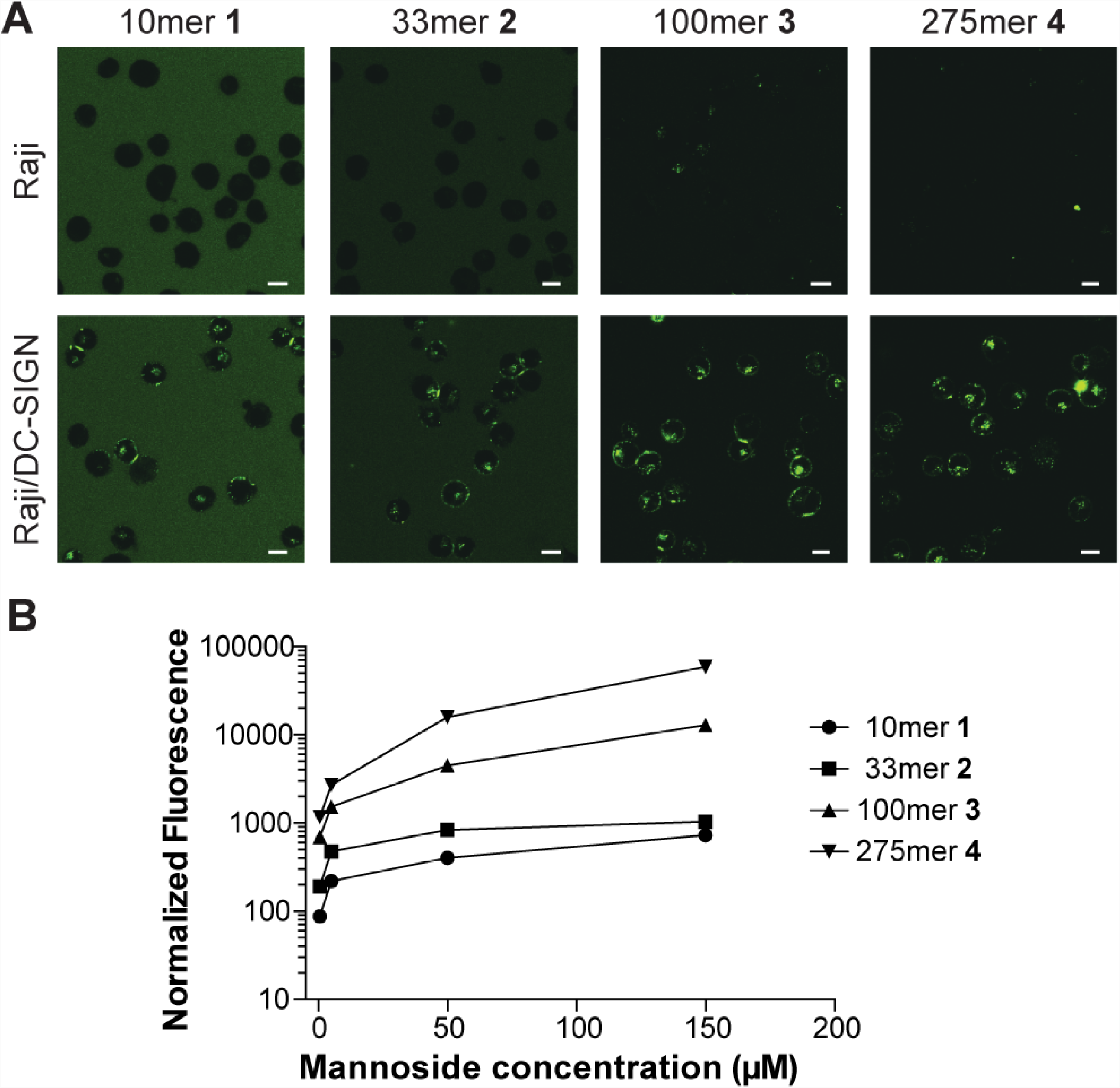
Comparison of fluorophore-labeled glycopolymer uptake in DC-SIGN-positive (Raji/DC-SIGN) and DC-SIGN-negative (Raji) cells. (A) Cells were treated with glycopolymers **1** – **4** (40 μM in mannoside ligand for **1** and **2**, 10 μM in mannoside ligand for **3** and **4**) for 30 min at 37 °C and internalization was visualized via confocal microscopy. Higher concentrations of glycopolymers **1** and **2** were required to visualize internalization. Scale bars, 10 μm. (B) Cells were treated with **1** – **4** (0.5, 5, 50, 150 μM in mannoside ligand) for 40 min, placed on ice, and flow cytometry was performed. The resulting fluorescence was normalized to account for the average number of fluorophores appended to each polymer.

To assess how the extent of DC-SIGN clustering influences internalization, Raji or Raji/DC-SIGN cells were treated with polymers of different lengths such that the mannose-ligand concentration was constant. The uptake of the longest polymer **4** (DP = 275) was approximately 10-fold higher than of the next longest **3** (DP = 100), which in turn was approximately 10-fold higher than for **2** (DP = 33) and **1** (DP = 10) (Fig. 2B). These findings indicate that receptor clustering indeed affects the efficiency of antigen internalization: the longer the polymer, the greater the extent of internalization.

### Endocytic trafficking of glycopolymer probes

Given that longer glycopolymers were more efficiently taken up than their shorter counterparts (Fig. 2), we assessed whether there were differences in glycopolymer trafficking. Mannosylated dendrimers had been shown to be internalized into endosomal compartments (33), and we assumed the glycopolymers would be routed to early endosomes. We therefore assessed whether the glycopolymers colocalized with transferrin, a marker for early endosomes. We employed Raji/DC-SIGN cells because they provide the means to dissect DC-SIGN’s role in the uptake of these antigens, as this receptor is the only cell surface C-type lectin expressed. To determine if uptake and trafficking is similar in DCs, we also tested monocyte-derived dendritic cells (moDCs). We found that each polymer colocalized with transferrin in either moDCs (Fig. 3B, C) or Raji/DC-SIGN cells (Fig. S2A,B). Trafficking occured via energy-dependent, DC-SIGN-mediated uptake (Fig. S2A-C). Treatment with anti-DC-SIGN antibodies abrogated uptake in Raji/DC-SIGN cells though inhibition of polymer uptake in moDCs was incomplete (Fig. S2C, D). Together, our data indicate that DC-SIGN mediates polymer internalization and trafficking, but suggest that other redundant receptors can effect uptake when DC-SIGN is inhibited. These data also show that polymer length modulates the efficiency of polymer internalization to endosomal compartments that lead to antigen processing.

**Fig. 3.**
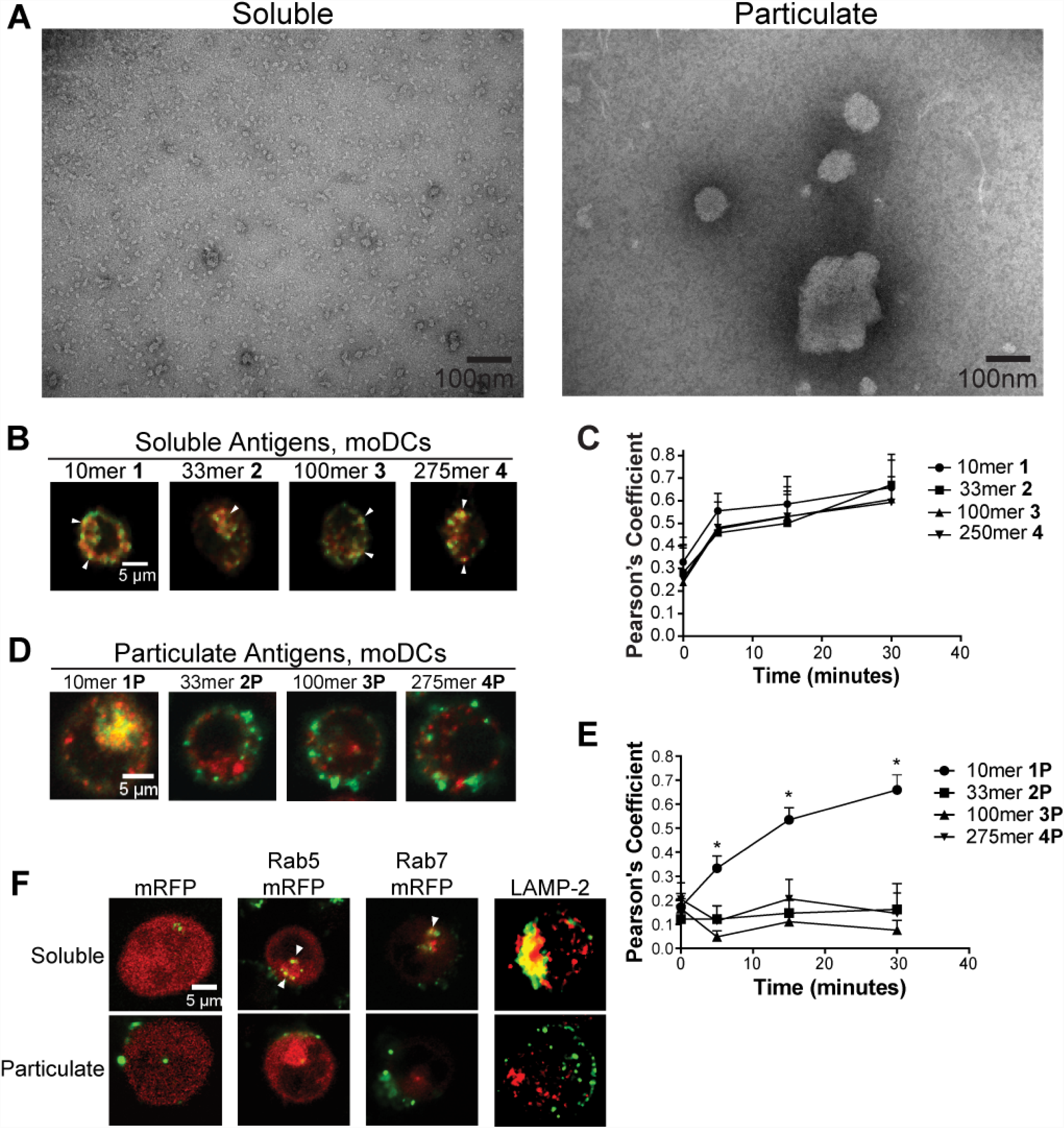
Trafficking of soluble and particulate glycopolymers. (A) Transmission electron microscopy of soluble and particulate 275mer polymers. (B) Soluble or (D) particulate glycopolymers (green) were added (mannose concentration of 10 μM) to moDCs at 37 °C for 30 min and their trafficking to transferrin-labeled early endosomes (red) was monitored via confocal microscopy. (C, E) Polymer/transferrin colocalization in panels B and D, respectively, was assessed at indicated time points. Pearson’s Coefficient was obtained by the Colocalization Threshold plugin in ImageJ using n>50 cells per treatment. Error bars represent the 95% confidence interval of the mean for representative data from at least 2 independent experiments. P values represent colocalization of the 10mer compared to each polymer at indicated time point. *p<0.001. (F) Polymer/Rab5, Rab7, or LAMP-2 colocalization was assessed in Raji/DC-SIGN cells either transfected with mRFP or Rab 5/7 mRFP or stained for LAMP-2 after treatment with polymer for 30 min at 37 °C. Scale bars are 100 nm in A and 5 μm in B-F.

### Differences in localization of particulate and soluble glycopolymers

The glycopolymers used in the aforementioned studies are soluble, but pathogens encountered by DCs (such as viruses and bacteria) are particulate. We therefore compared DC-SIGN-mediated internalization and trafficking for soluble versus particulate antigens. To access particulate antigens, we aggregated glycopolymers **1** – **4** by long term incubation of the compounds in solution. Analysis of the resulting polymers (termed **1p – 4p)** by dynamic light scattering (DLS) indicated that the length of the polymers influenced the particulate sizes that resulted. Specifically, glycopolymer **2p** (354 nm) afforded the smallest particles, and the longest polymers, **3p** and **4p**, gave rise to the largest particles (432 nm and 402 nm, respectively). Despite being exposed to conditions that led to the aggregation of the longer polymers, glycopolymer **1p** (10mer) did not aggregate, and its size was similar to that of the soluble polymers (1.4 nm) (Table S1). Additionally, application of transmission electron microscopy (TEM) to further characterize polymer size and structure revealed that both soluble and aggregate polymers were globular in shape but vastly different in size (Fig. 3A).

The trafficking of the particulate antigens differed markedly from that observed for the soluble polymers. Specifically, the particulate antigens were almost completely excluded from transferrin-labeled endosomes (Fig. 3D, E). Only glycopolymer **1p** colocalized with the endosomal marker transferrin, as it was similar in size to soluble glycopolymers **1** – **4** and exhibited the same trafficking patterns. In contrast, glycopolymers **2p – 4p** did not colocalize with transferrin; rather, they were localized to discrete puncta near the cell surface. We quantified the extent of colocalization over time in individual cells and observed that even at early timepoints, glycopolymers **2p** – **4p** were not trafficked to transferrin-positive endosomes. Thus, they appear to be directly routed to non-endosomal compartments. Additionally, while soluble polymers continued through the antigen presentation pathway, these aggregates failed to colocalize not only with transferrin but also with other endosomal markers, including Rab5 (a marker of early endosomes), Rab7 (a marker of late endosomes), and LAMP-2 (a marker of late endosomes/lysosomes) (Fig. 3F). Uptake and trafficking of the particulate antigens **2p – 4p** to these compartments occured via energy-dependent, DC-SIGN-mediated uptake (Fig. S3). The trafficking phenotype of particulate polymers is the same across Raji/DC-SIGN cells (Fig. S3C, D) and moDCs (Fig. 3D, E). Therefore, while polymer uptake is not completely dependent on DC-SIGN in moDCs, our findings indicate that DC-SIGN can mediate the non-endosomal trafficking in both cell types.

### Particulate antigens traffic to CD81^+^ compartments with HIV

The trafficking of the glycopolymer aggregates is reminiscent of the trafficking of HIV-1 in DCs. Only a subpopulation of HIV-1 internalized into DCs is routed to degradative compartments (34, 35). In contrast, HIV-1 is predominantly found in membrane-proximal, surface accessible invaginated pockets. These locations are identified by the presence of the tetraspanin CD81 (9). To test whether the internalized glycopolymers **2p – 4p** are trafficked to surface-accessible compartments, we added trypan blue, a dye that can quench AlexaFluor 488 emission and is not cell permeable (36, 37). The fluorescence emission of the particulate antigens was quenched, suggesting that these polymer aggregates are present in a cell-surface accessible compartment (Fig. 4A, B).

**Fig. 4.**
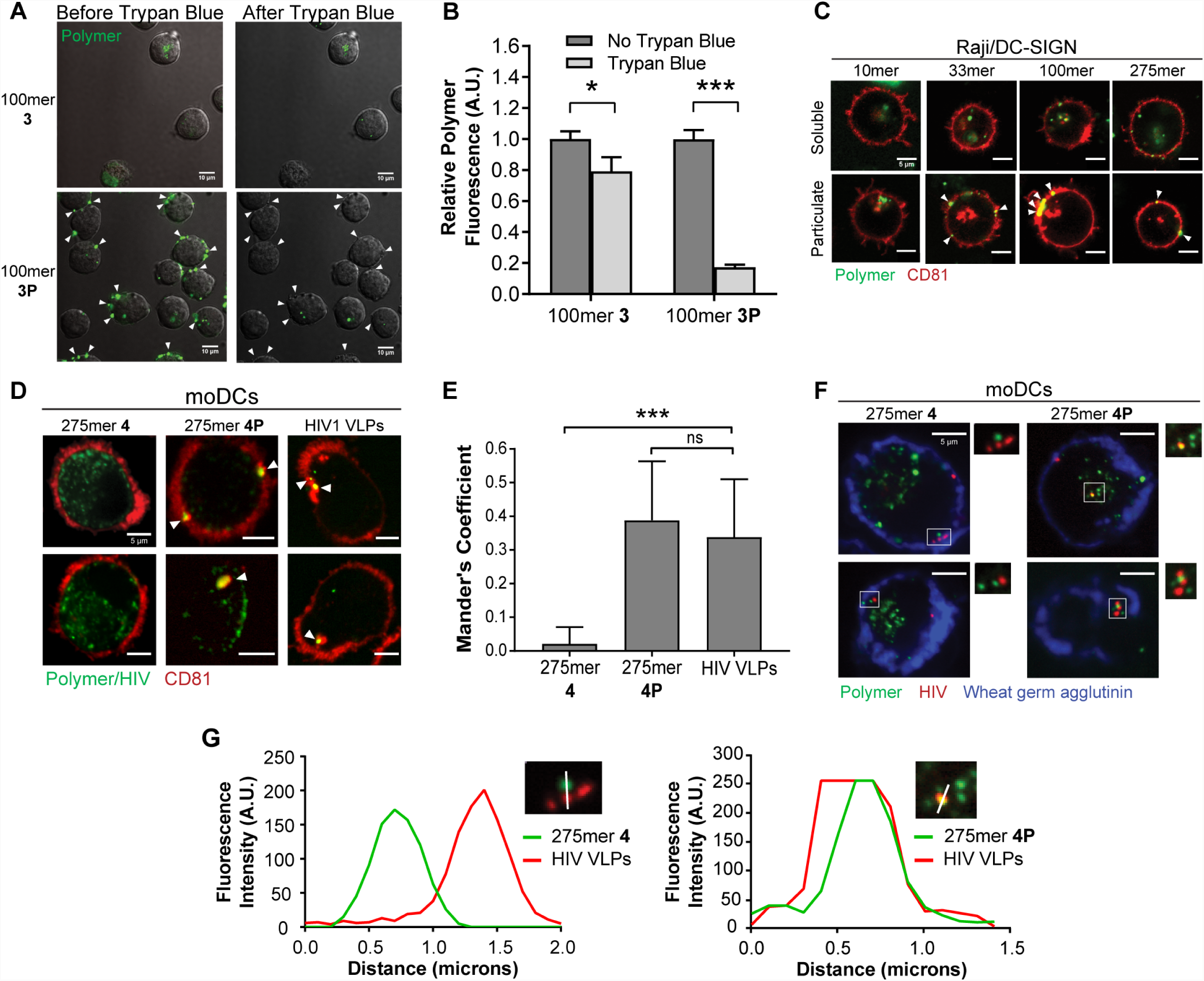
Comparison of trafficking of particulate and soluble antigens to HIV. (A) Soluble and particulate fluorophore-labeled glycopolymers (green, 10 μM mannose) were added to Raji/DC-SIGN cells at 37 °C for 30 min. Fluorescence quenching by trypan blue (2 min exposure) was monitored via confocal microscopy. Arrowheads indicate intracellular fluorescence not quenched by trypan blue. Scale bars, 10 μm. (B) Polymer fluorescence prior to and following trypan blue treatment was measured in ImageJ (normalized to untreated). Error bars represent the standard error of the mean of at least two experiments for n > 50 cells. (C, D) Polymers or fluorescently labeled HIV-1 VLPs (green) were added at 37°C to (C) Raji/DC-SIGN cells transfected with CD81-mCherry or (D) moDCs stained for endogenous CD81. Colocalization with CD81 was monitored by confocal microscopy. Arrowheads indicate regions of colocalization. (E) Quantitation of colocalization in moDCs in panel D, utilizing the Mander’s coefficient of antigen fluorescence overlapping with CD81. Error bars represent the standard error of the mean for at least two experiments for n **>**35 cells. (F) Polymer (green) and HIV (red) were coincubated in moDCs to assess colocalization. The plasma membrane was stained with wheat germ agglutinin (blue). (G) The overlap of polymer and HIV VLP fluorescence intensity (from F) was compared over the region indicated by a white line. Scale bars, 5 μm. *p<0.01, ***p<0.0001.

The trafficking data suggest that glycopolymer aggregates **2p** – **4p** are routed similarly to HIV-1. To evaluate this possibility, we examined whether **2p** – **4p** colocalized with CD81 in Raji/DC-SIGN cells (Fig. 4C). The soluble glycopolymers **1**– **4** did not colocalize with CD81. In contrast, CD81 localized with the fluorescence of particulate glycopolymers **2p** – **4p**. We next compared the routing of glycopolymers and fluorescently labeled HIV-1 virus like particles (VLPs) by analyzing their localization with CD81. VLPs resemble native virions and thus can be used to model viral uptake and trafficking from cell-to-cell at viral synapses (38–40). To circumvent potential effects of CD81-mCherry expression on polymer or HIV-1 uptake and trafficking, we examined antigen colocalization with endogenous CD81 in moDCs and Raji/DC-SIGN cells. While the soluble polymer **4** showed no appreciable colocalization, the aggregate **4p** and HIV-1 VLPs colocalized with dense pockets of CD81 in membrane proximal compartments (Fig. 4D, E and Fig. S4). To further validate this finding, we coincubated cells with HIV-1 VLP and polymer and found that aggregate **4p** directly colocalizes with HIV but soluble polymer **4** does not (Fig. 4F, G). These data confirm that aggregated polymers undergo the same route of internal trafficking as HIV-1.

## Discussion

We designed glycopolymer antigens to probe DC-SIGN endocytosis, and their application indicated that glycopolymer internalization depends upon its valency (length) — longer glycopolymers are internalized more readily than shorter glycopolymers. These results suggest that receptor clustering promotes DC-SIGN-mediated internalization. DC-SIGN is a tetramer with carbohydrate recognition domain (CRD) binding sites separated by about 40 Å (41, 42). Relatively small multivalent ligands may be able to cluster DC-SIGN and engage multiple CRDs for enhanced avidity (41, 43), but the shortest glycopolymer **1** (DP = 10) would likely only cluster two DC-SIGN tetramers at most. Glycopolymer **1**’s short (2.6 nm) length suggests it would not be capable of spanning the 4 nm distance between CRDs within a single DC-SIGN tetramer. That glycopolymer **1** is internalized supports previous findings that minimal DC-SIGN clustering can induce antigen internalization (44). Still, the uptake of this short polymer is significantly less efficient than that of the longer polymers, which can engage more binding sites and nucleate larger clusters. These data are consistent with a role for DC-SIGN clustering in antigen uptake.

Though the efficiency of uptake for the soluble glycopolymers varied with length, the trafficking of these mannosylated conjugates did not. All soluble mannosylated polymers were targeted to early endosomes. Since antigen routing to endosomes is an integral component of efficient antigen presentation in DCs (45-48), these findings suggest that soluble multivalent ligands can effectively deliver antigenic epitopes to the endosome for access to antigen presentation pathways. We anticipate that higher valency polymers may be more effective scaffolds for targeted synthetic vaccines than those glycoconjugates less capable of clustering DC-SIGN. This advantageous property of the higher valency polymers synergizes with their previously demonstrated ability to deliver multiple copies of antigenic peptides and therefore promote higher levels of T cell activation (28).

In striking contrast to the results with soluble polymers, the particulate antigens are almost entirely excluded from transferrin-labeled endosomes. Instead, particulate glycopolymers were routed to distinct CD81^+^ compartments and colocalized with HIV-1 particles. Another DC lectin, dectin-1 appears to make similar distinctions, as it preferentially internalizes particulate antigen over soluble antigen (49). Intriguingly, despite their distinct trafficking patterns, both small and large aggregates alike are internalized via energy dependent receptor-mediated uptake mechanisms. Antigen internalization depends only on DC-SIGN in Raji/DC-SIGN cells and partially on the lectin in moDCs. Consistent with these data, prior studies have shown that HIV internalization and transmission is inhibited by α-DC-SIGN blocking antibodies in Raji/DC-SIGN cells but not in DCs, implicating that other lectins are capable of facilitating viral uptake in when DC-SIGN is unavailable (50, 51). The polymer/HIV trafficking phenotypes observed in Raji/DC-SIGN cells are identical in moDCs, indicating that DC-SIGN can route antigen to distinct subcellular location based on the antigen’s structure (summarized in Fig. S6). Previous studies have demonstrated that changes in antigen size and shape can affect the extent of receptor-mediated internalization (reviewed in (52)), but to our knowledge, this is the first example where those features can direct antigen to be taken up by a receptor but trafficked to different compartments.

Super-resolution microscopy has shown that DC-SIGN exists on the cell surface in stable, loosely packed ∼80 nm diameter nanodomains that can be clustered into larger microdomains (53). These microdomains can range in size from 300 nm to 1.5 μm with an average spacing of 400 nm (54, 55). Based on their size, aggregate polymers can bridge several DC-SIGN nanodomains and even span multiple microdomains. These antigen attributes could potentiate unique internalization mechanisms. In immune cells, engagement and clustering of such receptor domains can lead to changes in actin organization (14-17, 56, 57). Recent studies have revealed that HIV-1 endocytosis into DCs and *trans* infection of CD4^+^ T cells at the viral synapse depends critically on actin remodeling factors, in particular dynamin-2 (10, 58). Similarly, the internalization of particulate antigens in our studies was partially dependent on dynamin-2 activity (Fig. S5). In contrast, the endocytosis of soluble antigens did not depend on dynamin-2 until later steps of trafficking. With these data, HIV-1 and particulate polymers may stimulate more profound changes in actin remodeling than soluble polymers, giving rise to their differential trafficking. The larger polymers may be capable of interfacing with and organizing DC-SIGN in a similar manner as HIV-1 resulting in alternative trafficking to the invaginated pockets. These findings suggest that the particulate polymer aggregates can be used to elucidate the mechanisms governing HIV’s divergent trafficking.

With the ability to control glycopolymer features, we found that the physical characteristics of a DC-SIGN-binding antigen affect its fate. In addition to the utility of polymers for understanding antigen uptake mechanisms, our findings provide opportunities to customize materials to serve as highly effective scaffolds for synthetic vaccines. By tailoring the properties of the polymer, one could direct antigenic cargo to specific compartments for classical presentation or cross presentation (46, 47). Our results also suggest strategies to target sites where viruses evade immune surveillance. In HIV-1 infected individuals, the persistence of HIV-1 in latently infected cells is a major obstacle toward completely eradicating the virus (59). Materials such as particulate glycopolymers could facilitate the identification of latently infected cells and potentially target and eliminate the viral reservoirs in these cells.

## Materials and Methods

### Polymer synthesis and characterization

A complete description of ligand and polymer synthesis and characterization is provided in the supporting information.

### Production of fluorescently labeled virus like particles

HEK293T cells plated in 10 cm tissue culture dishes were cotransfected to express plasmids encoding HIV-1 Gag-mCherry, HIV-1 Rev and HIV-1 CCR5 tropic Env. 48 hours post transfection, viral supernatants were collected, filtered through a 0.2 µm filter, and spun through a 0.1% sucrose cushion in a Beckman Coulter LE80K ultracentrifuge at 14000 RPM at 4 °C for 2 hours to sediment the virus like particles (VLPs). Pellet was re-suspended in 10 mL of culture media (DMEM 10% FBS 1% PenStrep 1% L-glutamine), filtered through a 0.2 µm sterile filter, aliquoted and stored at −80 °C until use.

### Cell Stimulation and Transfections

MoDCs, Raji or Raji/DC-SIGN cells were suspended at 1-4 x 10^6^ cells/mL in 1 mM CaCl_2_ 0.5 mM MgCl_2_ PBS pH 7.4 containing 0.1% BSA. Where indicated, cells were pre-treated at 37 °C for 15 min with inhibitors: Dynasore (80 μM), Genistein (100 μM) or vehicle (0.1% DMSO). With inhibitor present, cells were treated with indicated polymers at 10-40 µM mannose for indicated time periods prior to immunofluorescence analysis. For the expression of mCherry fusion constructs, Rab 5 and Rab 7 cDNA (a gift of M. Sandor) and human CD81 cDNA (Open Biosystems) were cloned into pmCherry-N1 vector (Clontech). Raji/DC-SIGN cells were electroporated with 1 µg of indicated plasmid using the Eppendorf Multiporator in iso-osmolar electroporation buffer using a 90 µs, 1200 V pulse. Cells were then transferred to pre-warmed culture media and utilized 48 hours post-transfection.

### Immunofluorescence and Confocal Microscopy

Where indicated, cells were incubated with 8 µg/mL Cy3-transferrin (Jackson Immunoresearch) for 30 min in 1% BSA/RPMI. Cells were washed and treated with polymer for the indicated time periods at either 4 °C or 37 °C and then placed on ice prior, washed and transferred to LabTek II Chambered Coverglass (#1.5). For HIV VLP internalization, cells were pre-treated overnight with LPS and then incubated with HIV-1::Gag-mCherry particles for 24 hr. In CD81 immunostaining experiments, cells were treated with polymer or HIV/VLP, fixed in 4 % paraformaldehyde, washed, permeabilized in 0.05 % saponin/PBS, and stained with mouse α-hCD81 (1.3.3.22) (Santa Cruz Biotechnology) and either Alexa Fluor 594-or Dylight 649-α-mouse IgG secondary antibody (Jackson Immunoresearch). The same procedure was used for LAMP2 colocalization experiments using rat α-LAMP2 primary antibody (Santa Cruz Biotechnology) and Alexa Fluor 594 donkey α-rat IgG secondary antibody (Thermo Fisher Scientific). For the trypan blue quenching experiments, fluorescence images were acquired before and after a 2 min incubation after trypan blue (0.1%; Bio-Rad), which was added carefully to minimize sample movement. Cells were imaged on a Nikon A1R confocal microscope (60X oil immersion lens, 1.4 NA) or an RPI spinning disk confocal microscope (63X oil immersion lens, 1.4 NA). Images were despeckled or background subtraction was performed prior to colocalization. Colocalization analysis was performed in Fiji using the Colocalization Threshold or Coloc 2 plugin to obtain the Pearson’s Coefficient and Mander’s coefficient of antigen overlap with transferrin, CD81, or LAMP2. All p-values reported were acquired from the Mann Whitney U-test performed using GraphPad Prism.

### Isolation and culture of monocyte-derived dendritic cells (moDCs)

Blood was drawn under UW Madison IRB approved Human Subjects minimal risk protocol M-2012-0130. Peripheral blood mononuclear cells from whole blood freshly drawn from healthy human volunteers were isolated via centrifugation using RosetteSep and Lymphoprep reagents (StemCell Technologies). Monocytes were separated by adherence to tissue culture flasks. Immature dendritic cells were prepared by treatment of monocytes with 500 U/ml human IL-4 and 800 U/ml human GM-CSF (R&D Systems) in CellGenix media for 6 days. Flow cytometry indicate that samples contain about 98% dendritic cells.

## Acknowledgements

This research was supported by the National Institutes of Health (AI055258). The authors thank the following people for assistance with transmission electron and confocal microscopy: Nikki Watson and Wendy Salmon of the W.M. Keck Biological Imaging Facility at the Whitehead Institute; Dr. Michelle Grevstad, of the UW Madison Biochemistry Optical Core; and Lance Rodenkirch, director of the UW Madison Keck Laboratory for Biological Imaging. Additionally, the authors acknowledge Dr. J. Hank, Dr. K. McDowell, and Prof. P. Sondel of University of Wisconsin – Madison for their assistance in obtaining blood.

