## Supporting Information for "Antigen structure affects cellular routing through DC-SIGN"

*General synthetic materials and methods.* All moisture- and oxygen-sensitive reactions were carried out in flame-dried glassware under a nitrogen atmosphere. Unless otherwise noted, all reagents and solvents were the highest commercially available grades and used without further purification. All chemicals were purchased from Sigma Aldrich with the exception of Alexa Fluor® 488 cadaverine sodium salt (Life Technologies). Dichloromethane ($CH_2Cl_2$) was distilled from calcium hydride, methanol (MeOH) was distilled from magnesium, and water ($H_2O$) was purified with a MilliQ purification system (Millipore). Analytical thin layer chromatography (TLC) was used to monitor reactions and was performed on 0.25 mm pre-coated Silica Gel 60 F254 (Merck). Compounds were visualized with ultraviolet light (254 nm) and/or charring with *p*-anisaldehyde (15 g *p*-anisaldehyde, 5 mL $H_2SO_4$, 1 mL AcOH, 250 mL ethanol). Flash chromatography was performed on 230 – 400 mesh SiliaFlash® P60 silica gel (Silicycle).

$^{1}H$ and $^{13}C$ nuclear magnetic resonance (NMR) spectra were recorded on Bruker AC-300 or Varian MercuryPlus 300 spectrometers, and polymers $^{1}H$ NMR spectra were recorded Varian INOVA 600 spectrometer. Chemical shifts were reported relative to trimethylsilane or residual solvent peaks in parts per million ($CHCl_3$: $^{1}H$ δ 7.26, $^{13}C$ δ 77.0; $CH_3OH$: $^{1}H$ δ 3.31, $^{13}C$ δ 49.0; DMSO-$d_6$: $^{1}H$ NMR δ 2.50). Peak multiplicity is reported as singlet (s), doublet (d), multiplet (m), doublet of doublet (dd), etc. All spectra are reported in the supplemental information. High resolution electrospray ionization mass spectra (HRESI-MS) were obtained on a Micromass LCT (electrospray ionization, time-of-flight analyzer).

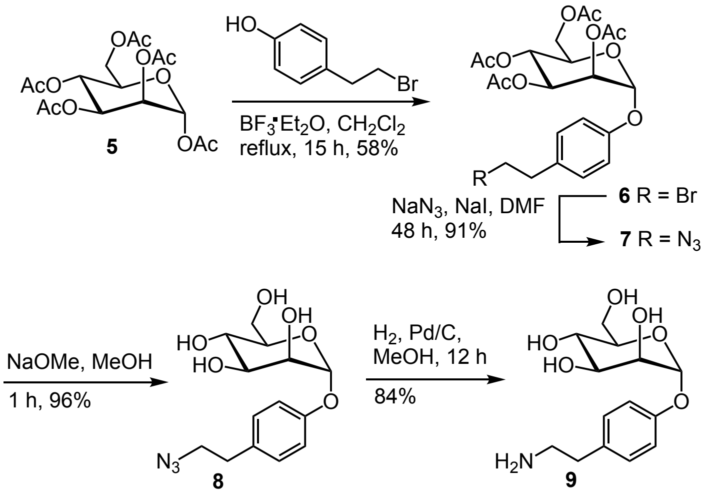

**Scheme S1.** Synthetic route for mannoside group appended to polymers.

*α-1-O-(4-hydroxyphenylethylbromide)-2,3,4,6-tetra-O-acetyl-D-mannoside **6***. To a flask containing peracetylated mannoside **5** (0.31 g, 0. 80 mmol) in $CH_2Cl_2$ (4 mL), 4 Å molecular sieves were added and stirred for 30 min. The solution was cooled to 0°C, and 4-hydroxyphenylethyl bromide (0.48 g, 2.4 mmol) was added. Boron trifluoride diethyl etherate (0.39 mL, 3.2 mmol) was added dropwise. The solution was maintained at reflux overnight. The reaction mixture was quenched with triethyl amine. The organic layer was washed with saturated $NaHCO_3$ and brine, dried over $MgSO_4$, filtered, and concentrated under reduced pressure. The crude oil was purified by column chromatography (3:7 EtOAc:hexanes) to afford 12 (0.25 g, 0.46 mmol, 58%) as a colorless oil. $^1$H NMR (300 MHz, $CDCl_3$) δ 7.21 – 7.11 (m, 2H), 7.09 – 6.98 (m, 2H), 5.56 (dd, *J* = 10.0, 3.5 Hz, 1H), 5.51 (d, *J* = 1.8 Hz, 1H), 5.43 (dd, *J* = 3.5, 1.8 Hz, 1H), 5.37 (t, *J* = 10.1 Hz, 1H), 4.35 – 4.22 (m, 1H), 4.16 – 4.02 (m, 2H), 3.53 (t, *J* = 7.5 Hz, 2H), 3.11 (t, *J* = 7.5 Hz, 2H), 2.20 (s, 3H), 2.05 (s, 9H). $^{13}$C NMR (75 MHz, $CDCl_3$) δ 170.48, 169.93, 169.90, 169.70, 154.59, 133.53, 129.83, 116.64, 95.90, 69.42, 69.15, 68.88, 65.97, 62.12, 38.49, 33.04, 20.68. HRMS (EMM): calc'd for $C_{22}H_{27}BrO_7$ $[M+Na]^+$: 553.0680, found 553.0699.

*α-d-1-O-(4-hydroxyphenylethylazide)-2,3,4,6-tetra-O-acetyl-D-mannoside* **7**. To a flask containing **6** (2.39 g, 4.49 mmol), $NaN_3$ (0.880 g, 13.5 mmol) and NaI (0.670 g, 13.5 mmol) were added. DMF (15 mL) was added, and the solution was stirred for 48 h. $CH_2Cl_2$ (60 mL) was added, and the solution was washed with water (20 mL) and brine (20 mL). The organic layer was dried with $MgSO_4$, filtered, and concentrated under reduced pressure to afford **7** (1.98 g, 4.0 mmol, 91%) as a pure yellow solid. $^1$H NMR (300 MHz, $CDCl_3$) δ 7.19 – 7.12 (m, 2H), 7.07 – 7.01 (m, 2H), 5.56 (dd, *J* = 10.0, 3.5 Hz, 1H), 5.50 (d, *J* = 1.9 Hz, 1H), 5.44 (dd, *J* = 3.5, 1.8 Hz, 1H), 5.37 (t, *J* = 10.2 Hz, 1H), 4.28 (dd, *J* = 12.1, 5.2 Hz, 1H), 4.08 (ddd, *J* = 12.5, 8.4, 3.6 Hz, 2H), 3.47 (t, *J* = 7.1 Hz, 2H), 2.84 (t, *J* = 7.1 Hz, 2H), 2.19 (d, *J* = 0.6 Hz, 3H), 2.05 (s, 3H), 2.03 (s, 3H). $^{13}$C NMR (75 MHz, $CDCl_3$) δ 170.46, 169.89, 154.50, 132.68, 129.91, 116.72, 95.91, 69.41, 69.14, 68.89, 65.96, 62.12, 52.49, 34.51, 20.85. HRMS (EMM) calc'd for $C_{22}H_{27}N_3O_{10}$ $[M+NH_4]^+$: 511.2035, found 511.2015.

*α-1-O-(4-hydroxyphenylethylazide)-D-mannoside **8***. To a solution of **7** (0.140 g, 0.284 mmol) in methanol (2.9 mL), 0.5 M sodium methoxide in methanol (0.60 mL, 0.28 mmol) was added. The solution was stirred for 1 h, and then quenched with strongly acidic resin. The solution was filtered and concentrated under reduced pressure. The oil was purified by column chromatography (3:14 MeOH:$CH_2Cl_2$) to afford **8** (0.089 g, 0.27 mmol, 96%) as a colorless residue. $^1H$ NMR (300 MHz, $CD_3COD$) δ 7.15 (d, $J$ = 8.6 Hz, 2H), 7.07 – 7.02 (m, 2H), 5.47 (d, $J$ = 1.7 Hz, 1H), 4.03 (dd, $J$ = 3.3, 1.8 Hz, 1H), 3.94 (dd, $J$ = 9.4, 3.3 Hz, 1H), 3.85 – 3.71 (m, 3H), 3.61 (dt, $J$ = 9.8, 3.7 Hz, 1H), 3.42 (t, $J$ = 7.0 Hz, 2H), 2.78 (t, $J$ = 7.1 Hz, 2H). $^{13}C$ NMR (75 MHz, $CD_3OD$) δ 155.51, 132.45, 129.80, 116.80, 99.03, 74.03, 71.30, 70.84, 61.42, 52.49, 34.28. HRMS (EMM) calc'd for $C_{14}H_{19}N_3O_6$ $[M+Na]^+$: 348.1167, found 348.1150.

*α-1-O-(4-hydroxyphenylethylamino)-D-mannoside **9***. To a solution of **8** (0.089 g, 0.27 mmol) in methanol (1.4 mL), palladium on carbon (0.017 g) was added. The flask was evacuated under a balloon of hydrogen gas and left overnight. The mixture was filtered through a Celite plug, and the plug was washed with methanol. The solution was concentrated in vacuo to afford **9** (0.063 g, 0.23 mmol, 84%) as a clear residue. The product purity was verified via $^1H$ NMR. $^1H$ NMR (300 MHz, $CD_3OD$) δ 7.18 – 7.09 (m), 7.09 – 6.97 (m), 5.44 (d, $J$ = 1.8 Hz), 4.01 (ddd, $J$ = 5.3, 3.4, 1.9 Hz), 3.92 (ddd, $J$ = 9.0, 5.3, 3.3 Hz), 3.81 – 3.70 (m), 3.61 (ddt, $J$ = 9.7, 4.3, 2.7 Hz), 3.35, 2.86 – 2.74 (m), 2.69 (q, $J$ = 6.6 Hz). $^{13}C$ NMR (75 MHz, $CD_3OD$) δ 156.57, 134.98, 130.91, 118.06, 100.44, 75.39, 72.55, 72.15, 68.46, 62.75, 51.88, 44.43, 39.48, 35.78. HRMS (EMM) calc'd for $C_{14}H_{21}NO_6$ $[M+H]^+$: 300.1442, found 300.1145.

*5-oxo-norbornene 2-exo-N-hydroxylsuccinimidyl ester **10***. To a flask containing exo-5-*oxo*-norbornene-2-carboxcylic acid (1) (0.048 g, 0.35 mmol) in dichloromethane (1 mL), *N*-hydroxysuccinimide (0.053 g, 0.49 mmol) and 1-ethyl-3-(3-dimethylaminopropyl)carbodiimide (0.081 g, 0.45 mmol) were added under argon. The solution was stirred for 14 h and then concentrated under reduced pressure. The residue was purified by column chromatography (1:4 to 1:1 EtOAc:hexanes) to afford monomer **10** (0.028 g, 0.14 mmol, 37%) as a white solid. $^1H$ NMR (300 MHz, $CDCl_3$) δ 6.47 (dd, $J$ = 5.8, 1.7 Hz, 1H), 6.40 (dd, $J$ = 5.8, 1.7 Hz, 1H), 5.34 (t, $J$ = 1.2 Hz, 1H), 5.19 – 5.11 (m, 1H), 2.92 – 2.78 (m, 4H), 2.72 (dd, $J$ = 8.5, 4.0 Hz, 1H), 2.25 (dt, $J$ = 11.6, 4.3 Hz, 1H), 1.72 (dd, $J$ = 11.7, 8.5 Hz, 1H).

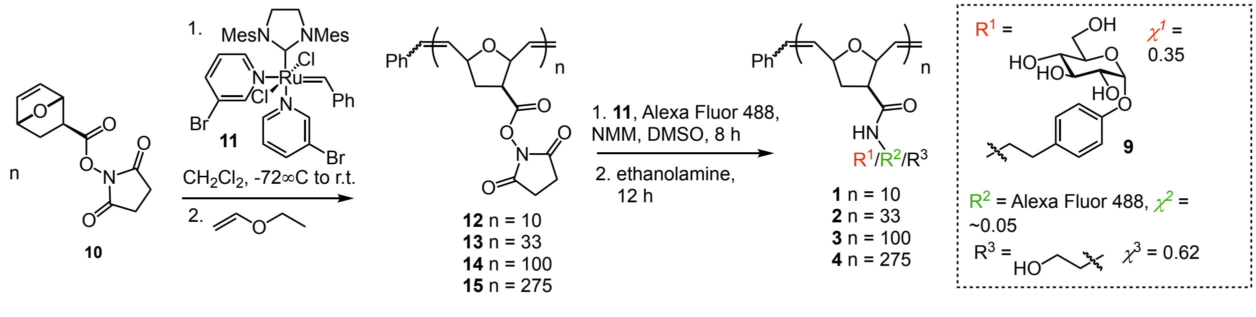

**Scheme S2. Synthetic route to glycopolymers using ROMP.**

**General polymer synthesis**. The synthesis of 10-mer **12** is provided as a general protocol. The degree of polymerization was adjusted by varying the catalyst-to-monomer ratio. To a vial containing monomer **10** (0.098 g, 0.42 mmol), $CH_2Cl_2$ (4.1 mL) was added, and the solution was cooled to -72°C under argon. A solution of catalyst **11** in $CH_2Cl_2$ (0.1 *M*, 0.42 mL, 0.042 mmol) was added. The solution was allowed to warm to 0 °C, at which point TLC analysis indicated no monomer remained. Ethyl vinyl ether (0.2 mL, 2.0 mmol) was added, and the solution was stirred overnight at rt. The polymer was precipitated into ether (45 mL), and the suspension was centrifuged at 3,000 rpm for 20 min. The supernatant was decanted, and this procedure was repeated two more times. The polymer was allowed to dry overnight. Comparison of the integration values of the phenyl end cap moiety to the vinylic protons of the polymer backbone by $^1H$ NMR spectroscopy gave the degree of polymerization. Polydispersity indices (PDIs) and number-average molecular weights ($M_n$) were measured on a Waters gel permeation chromatography (GPC) using two Viscotek ViscoGEL I-series columns (I-MBL MW-3078 and I_MBH MW-3078) with 0.1 M LiBr in DMF as the eluent, and Varian polyethylene glycol/polyethylene oxide polymer were used as standards.

**Polymer functionalization**. The functionalization procedure for 10-mer **12** is provided. The mannose epitope to polymer molar ratio could be varied to alter polymer density. Polymer **3** (1.97 mg, 0.00080 mmol) was dissolved in DMSO (0.8 mL) in a polypropylene tube. A solution of **9** in DMSO (100 mg/mL, 9.6 µL, 0.0032 mmol) was added followed by a solution of Alexa Fluor® 488 cadaverine sodium salt in DMSO (10 mg/mL, 77 µL,1.2 µmol). The tube was protected from light and spun gently overnight. Ethanolamine (1.2 µL, 0.02 mmol) was added and the resulting solution was spun gently on rotisserie for 8 h. The polymer solution was transferred to 2K MWCO dialysis tubing and dialyzed three times against water. The water was removed by lyophilization to afford glycopolymer **1** as an orange powder. Percent conjugation of the mannoside epitope was calculated by $^1H$ NMR spectroscopy. For the longer polymers, aggregates formed over time (several months) when left at room temperature in a concentrated DMSO solution. The speed of aggregate formation could be enhanced by incubation in a high salt buffer (1.8M phosphate buffer, pH 7.3) at room temperature. Aggregate size was consistent using both aggregation methods.

**Dynamic light scattering.** Dynamic light scattering was performed on a DynaPro NanoStar DLS. All polymer solutions were measured at 40 µM in phosphate buffered saline (pH 7.4). DLS measurements were performed in triplicate and evaluated with regularization analysis. The size distributions are provided in Table S1 below.

**Transmission electron microscopy.** Polymer was diluted to 20 μM in phosphate-buffered saline (pH 7.4). Carbon coated grids were ionized and floated in polymer for 1 minute. The grid was washed with tris buffer and floated on 1-2% acidic uranyl acetate for 20-60 seconds. Excess stain was drawn off with filter paper, and the grid was air dried. Polymer was then imaged using a FEI Technai Spirit Transmission Electron Microscope.

**Cell culture**. Raji and Raji/DC-SIGN cells were obtained from the NIH AIDS Reference and Reagent Program (2). Cells were maintained in RPMI media (Gibco) supplemented with 10% fetal bovine serum and penicillin/streptomycin at 37 °C and 5% carbon dioxide.

**Polymer internalization confocal microscopy.** Raji and Raji/DC-SIGN cells were spun and resuspended in PBS supplemented with $CaCl_2$ (Gibco) and 1% BSA at one million cells per mL. Polymers were added to 200,000 cells to a final mannose concentration of 40 μM (10-mer and 33-mer) or 10 μM (100-mer and 275-mer). Cells were then incubated with polymer for 30 minutes at 37°C, washed, and transferred to eight-chambered cover glass slides (Nunc). Cells were visualized on a Nikon Eclipse Ti-E A1R confocal microscope with a 60x oil immersion lens and the NIS Elements software package.

**Flow cytometry.** Raji and Raji/DC-SIGN cells were treated with each polymer at mannose residue concentrations that ranged from 0.5 to 150 μM. Polymer interaction with cells was analyzed using a BD FACSCalibur flow cytometer. Geometric mean of AF488 fluorescence for each sample was calculated using the FlowJo software package. To identify DC-SIGN-specific polymer interactions, the fluorescence value measured for Raji cells was subtracted from that of Raji/DC-SIGN cells for each point measured. These values were then normalized to fluorophore content per polymer. In Figure 2, the fluorescence values measured at 0.5 μM mannose were set to one for each curve. For receptor blocking antibodies studies, cells were pre-treated with antibodies against the indicated receptor (α-DC-SIGN [BD Bioscience, 10 μg/mL], α-Dectin-2 [R&D Systems, 20 μg/mL], α-CD206 [BD Biosciences, 20 μg/mL]) for 20 minutes on ice, immediately followed by incubation with 0.5 μM polymer for 15 minutes at 37 °C.

**Table S1.** Polymer sizes as determined by dynamic light scattering (DLS)..

| Glycopolymer | Diameter (nm) |
|---|---|
| 1 | 2.6 |
| 2 | 2.0 |
| 3 | 9.0 |
| 4 | 8.8 |
| 1p | 1.4 |
| 2p | 354 |
| 3p | 432 |
| 4p | 402 |

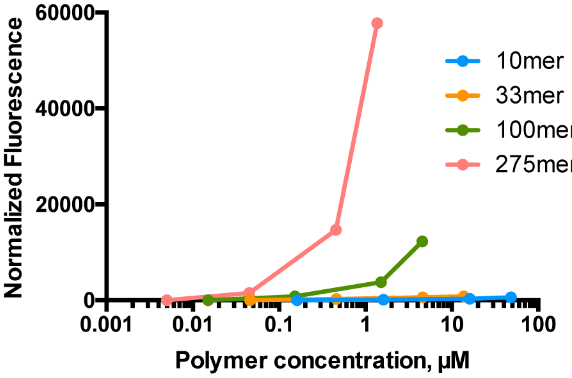

**Fig. S1.** Dose dependence of polymer internalization. Raji and Raji/DC-SIGN cells were treated with glycopolymers **1** – **4** at a range of concentrations for 30 min at 37 °C. Samples were placed on ice and fluorescence was measured via flow cytometry. Normalized fluorescence was calculated by subtracting Raji fluorescence signal from Raji/DC-SIGN fluorescence signal and normalizing to the number of fluorophores per polymer.

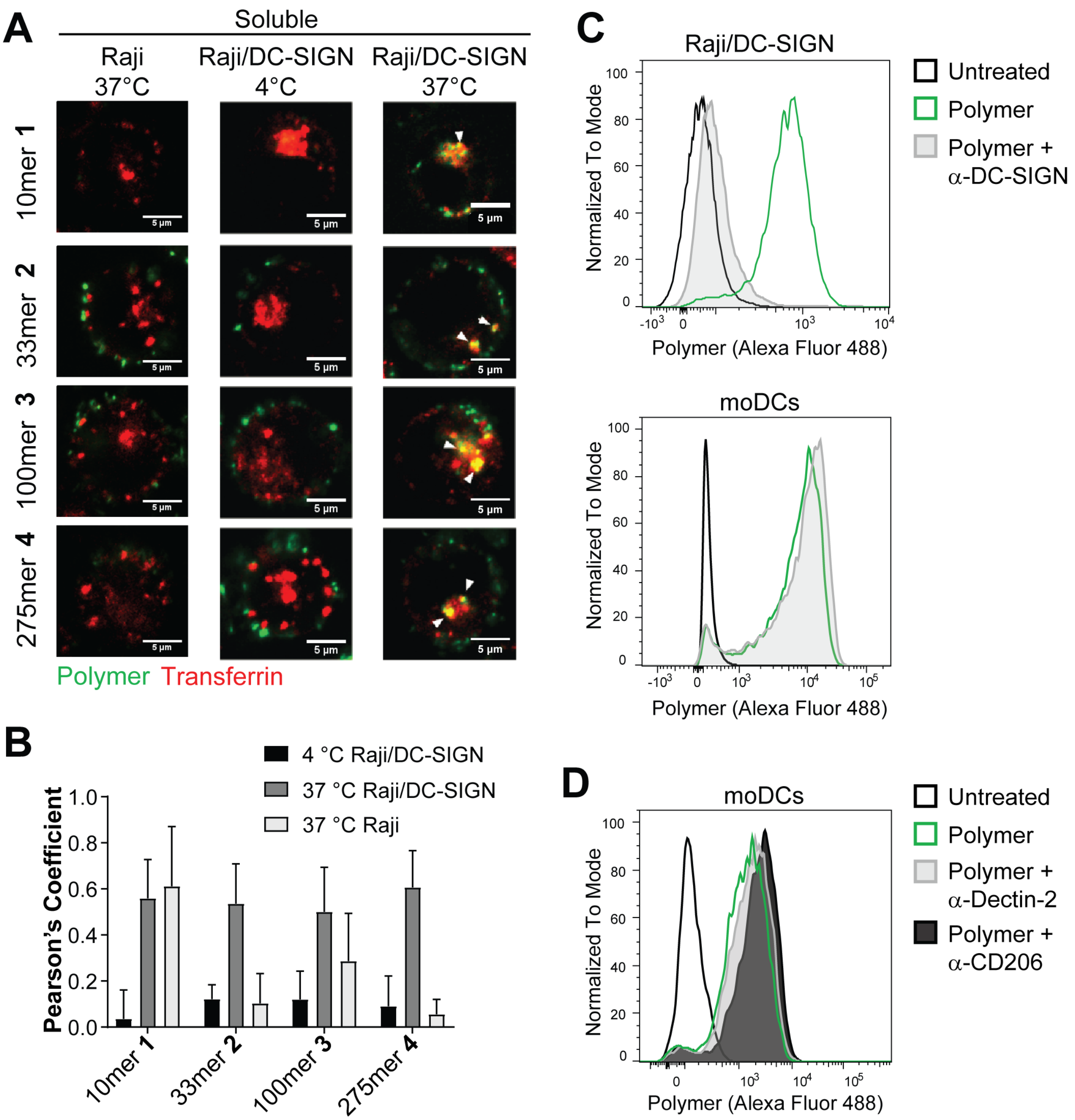

**Fig. S2.** DC-SIGN-mediated internalization and trafficking of soluble (**1**–**4**) polymers. (A) Soluble glycopolymers (green) were added (10 µM mannose) to Raji or Raji/DC-SIGN cells at 4 °C or 37 °C for 30 min. Trafficking to transferrin-labeled early endosomes (red) was monitored via confocal microscopy. Scale bars, 5µm. (B) The colocalization of each polymer with transferrin was assessed for n ≥ 10 cells per treatment using the Pearson's Coefficient obtained by the Colocalization Threshold plugin in ImageJ. (C) Raji/DC-SIGN or moDCs were pre-treated with α-DC-SIGN blocking antibody, followed by polymer (100mer **3**) to assess DC-SIGN specific uptake. (D) moDCs were treated with α-Dectin-2 or α-CD206 blocking antibody and polymer (100mer **3**) to assess uptake by the given receptor.

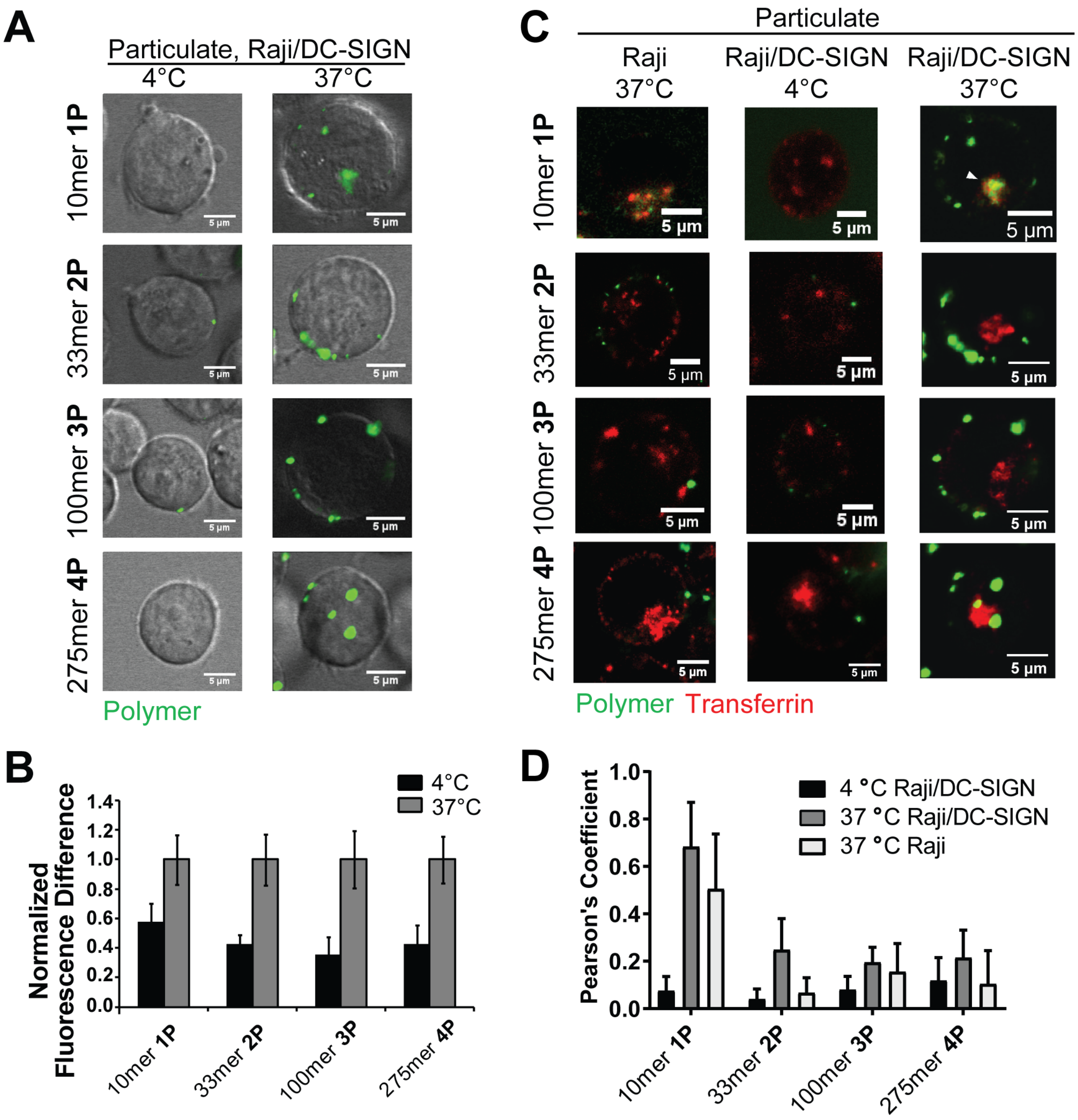

**Fig. S3.** DC-SIGN-mediated internalization and trafficking of particulate (**1P**–**4P**) polymers. (A) Internalization of particulate polymers (10 µM mannose residue concentration) was assessed in Raji/DC-SIGN cells at 4 °C or 37 °C. (B) Total cell associated polymer fluorescence intensity from A was measured in ImageJ for n>15 cells. Data are normalized to the 37°C polymer treatment conditions for each polymer. (C) Particulate glycopolymers (green) were added (10 µM mannose) to Raji or Raji/DC-SIGN cells at 4 °C or 37 °C for 30 min. Trafficking to transferrin-labeled early endosomes (red) was monitored via confocal microscopy. (D) The colocalization of each polymer with transferrin was assessed for n ≥ 10 cells per treatment using the Pearson's Coefficient obtained by the Colocalization Threshold plugin in ImageJ. Scale bars, 5µm.

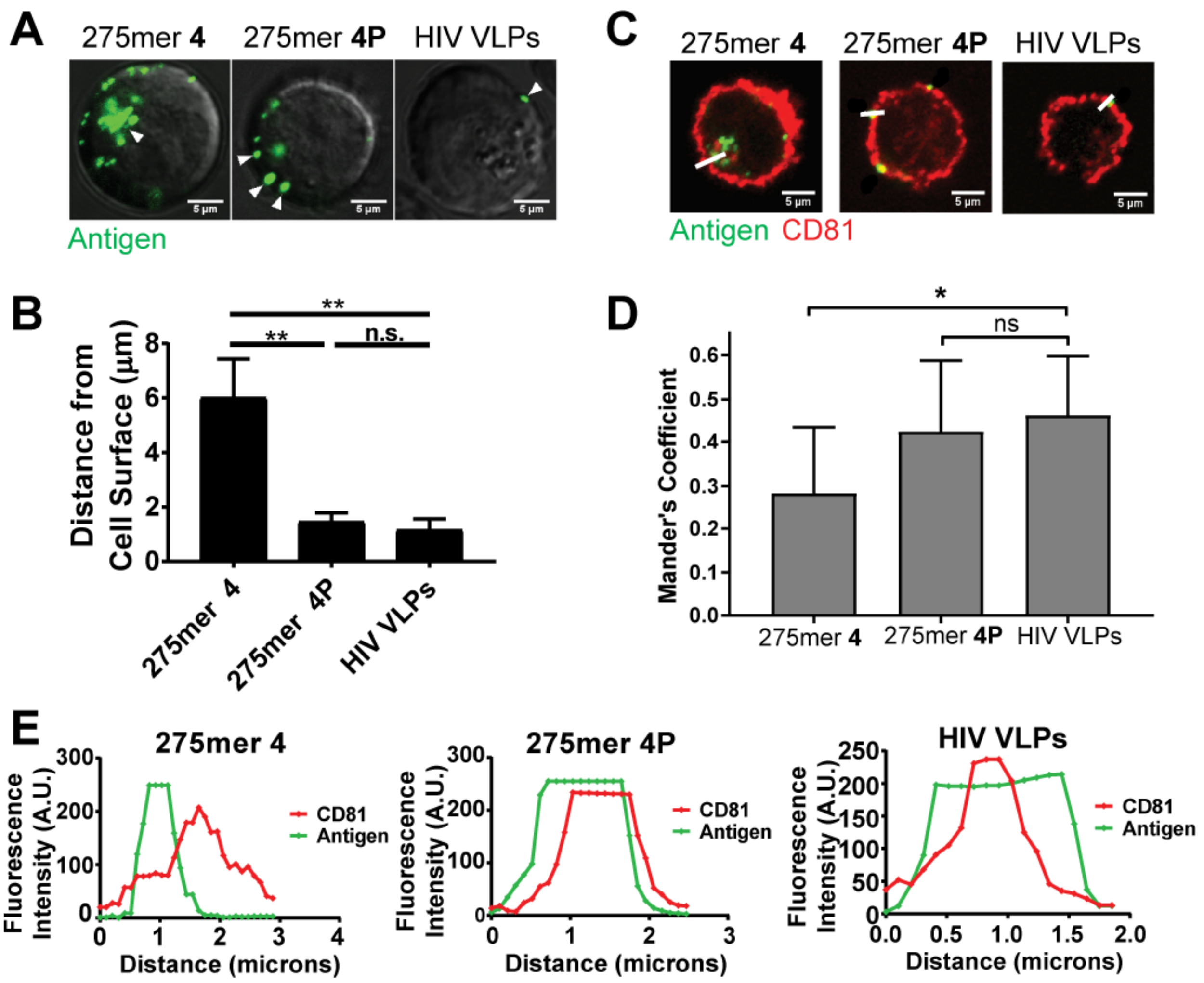

**Fig. S4.** Localization of particulate polymers and HIV-1 VLPs in Raji/DC-SIGN cells. (A) The proximity of particulate polymers (10 µM mannose) or HIV VLPs to the cell surface was assessed in Raji/DC-SIGN cells after 30 min treatment at 37 ºC. (B) Distance of polymer staining to the cell surface from (A) was measured in ImageJ for n > 5 cells. (C-E) In Raji/DC-SIGN cells treated with HIV-1 VLPs or polymers (C), (the colocalization with CD81 was assessed for n>15 cells (D), and the overlap of polymer or HIV-1 fluorescence intensity with CD81 fluorescence was compared over the region indicated by a line in (E). Scale bars, 5 µm. *p<0.001, **p<0.0005

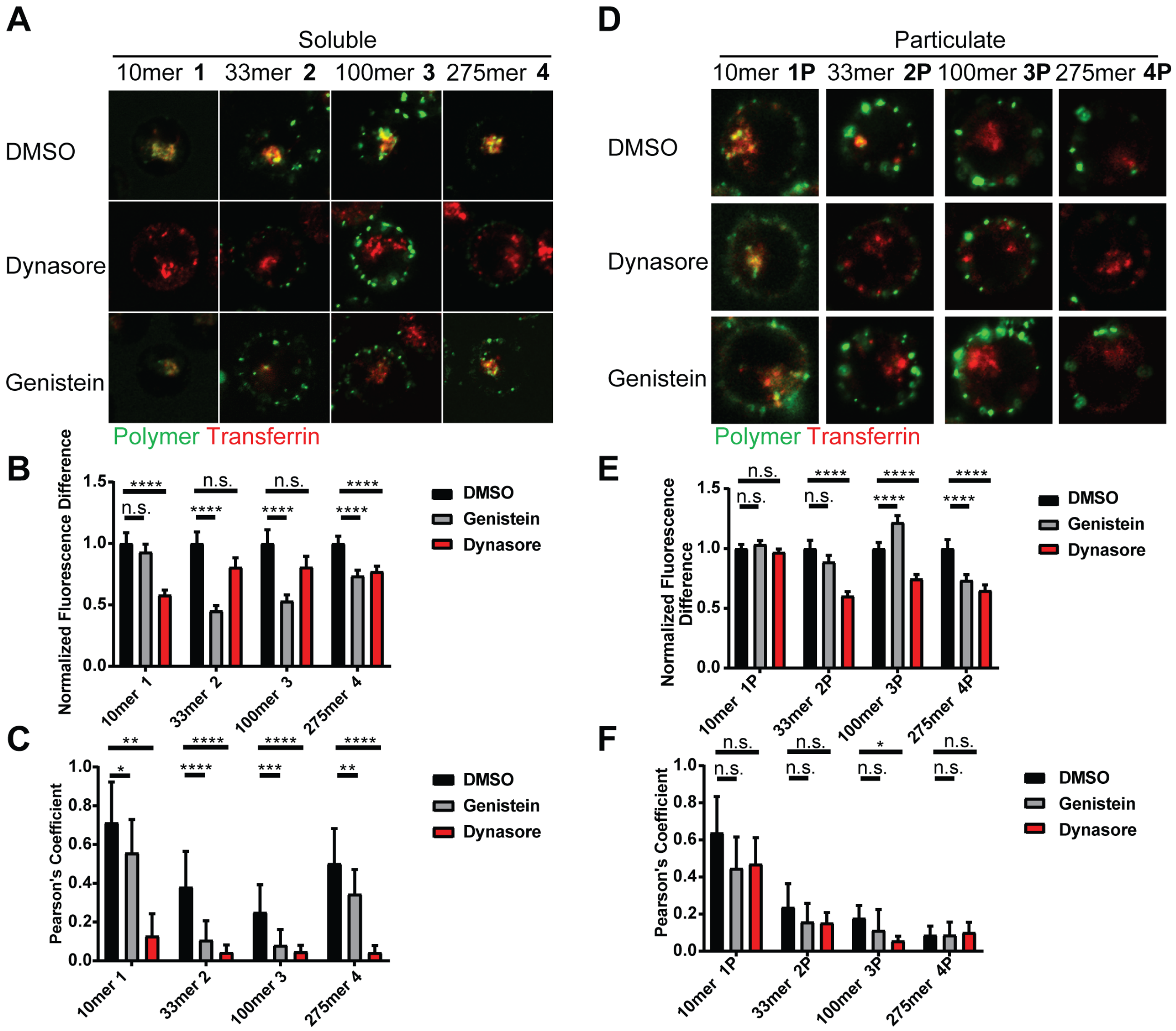

**Fig. S5.** Probes of endocytic mechanisms of glycopolymer trafficking. (A, D) After 10 min pre-treatment with 10 μM dynasore, 20 μM genistein or vehicle, (A) soluble or (D) particulate glycopolymers (green) were added (mannose concentration of 10 μM) to Raji/DC-SIGN cells at 37 °C for 30 min. Trafficking to transferrin-labeled early endosomes (red) was monitored via confocal microscopy. (B, E) The effect of endocytic inhibitors on polymer internalization in (A) and (D) respectively was assessed. Total cell-associated polymer fluorescence intensity was measured in ImageJ for n >15 cells. Data are normalized to the DMSO treatment condition for each polymer. (C, F) Polymer colocalization with transferrin in (A) and (D) was quantified in using the Pearson's Coefficient obtained by the Colocalization Threshold plugin in ImageJ. Error bars represent the standard deviation for n > 15 cells. $*p < 0.05$, $**p < 0.01$, $***\ p < 0.001$, $****\ p < 0.0001$.

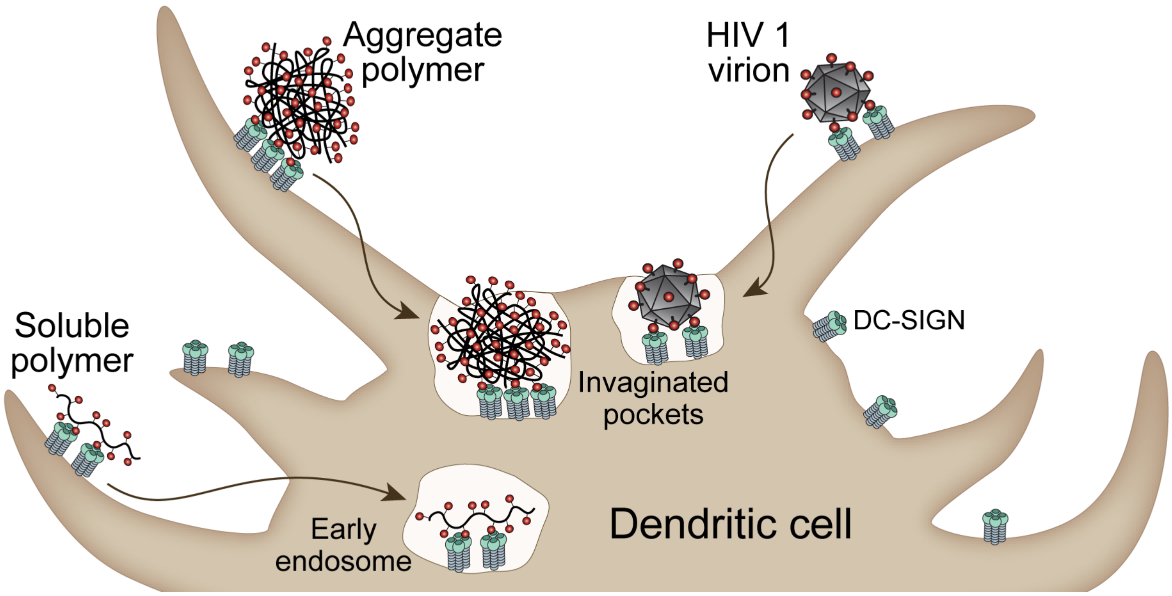

**Fig. S6**. Summary of differential routing of soluble and particulate polymers by DC-SIGN. Soluble polymers are internalized and routed toward early endosomes. Aggregated polymers are internalized to non-endosomal invaginated pockets and co-localize with HIV-1 virions.
